# Ecological background predicts object feature salience in *Sylvaemus uralensis* and *Apodemus agrarius*

**DOI:** 10.64898/2026.04.08.717221

**Authors:** M. Yurin Alexander, A. Solodova Ekaterina, A. Egovtsev Nikita, M. Malygin Vasily, Viktor Yu. Oleinichenko, G. Pleskacheva Marina

## Abstract

Animals should attend preferentially to stimulus features relevant to their ecological niche, but whether this predicts which object features drive exploration in behavioral assays has not been, to our knowledge, systematically tested. We presented two wild-caught mouse species with contrasting ecologies – wood mice (*Sylvaemus uralensis*), which inhabit forests and regularly climb, and striped field mice (*Apodemus agrarius*), which occupy open habitats – with six pairs of objects assembled from standardized components, the members of each pair differing along exactly one researcher-defined feature: height, colour, apex shape, aperture presence, or decorative motif. Height produced a species-specific, ecology-predicted response: *S. uralensis* avoided the shortest object, converging across zone occupancy, tactile contact, and climbing, while *A. agrarius* showed no comparable avoidance. No other feature produced a detectable preference in either species. Two of these nulls were features we had also expected to be salient on general ecological grounds, though without predicting a species difference; only the comparative prediction, height, was borne out. General biological plausibility of this kind, reasoned by an experimenter, is therefore not sufficient to establish that a feature is salient, and paired-object testing against single features offers a tractable way to find out which are, in a given species, before those objects are used in memory or novelty assays.

## Introduction

Animals prioritize ecologically relevant stimulus dimensions over arbitrary physical features when evaluating their environment. This principle, well established in the comparative cognition literature (Shettleworth, 2010)[1], predicts that the features animals attend to when encountering unfamiliar objects will reflect their species-typical ecology and behavioral repertoire rather than the perceptual dimensions a human observer might consider salient. If feature salience is species-specific in this way, then the features that drive differential exploration of objects in controlled behavioral assays should differ predictably between species with contrasting ecologies. This prediction, to our knowledge, has not been systematically tested in object exploration assays.

Solving this problem has both basic and applied implications. On the basic side, testing feature salience may help us in evaluating natural animal preference for individual features and their relative importance in laboratory and natural contexts. That in turn may inform us of interspecies– or interstrain-specific behavioral differences in cognition that are not obviously extractable from existing task paradigms. On the applied side, object-based behavioral tasks are central to cognitive research in rodents (Leger *et al.*, 2013; Lueptow, 2017)[2,3], and whether animals discriminate between objects on the basis of arbitrary perceptual features, ecological valence, or some combination shapes how the results of such tasks should be interpreted. Object selection across laboratories remains largely unsystematic, and the assumptions underlying object-based tasks have themselves been questioned: Ennaceur (2010)[4] noted that work in rats and mice has focused on memory while neglecting object perception and the features animals actually represent and a survey of 69 mouse NOR reports published in Nature-family journals during 2024 found that only 24.6% described the objects used and only 14.5% employed a cross-over design in which the same object served as familiar for one group and novel for another (Koivisto *et al.*, 2025)[5]. Such controls matter: laboratory mice preferentially explore objects made of shiny materials such as metal and glass, and also larger objects offering greater affordance (Tropea *et al.*, 2022)[6], and exploration levels and the actions animals direct at an object depend on its functional properties rather than its appearance (Heyser and Chemero, 2012)[7]. Existing recommendations address only whether two complete objects produce equal exploration, not which specific features drive any observed preference (Leger *et al.*, 2013; Lueptow, 2017)[2,3], and recent efforts toward physical standardization through 3D-printed object libraries (Inayat *et al.*, 2021)[8] leave open the question of which features within those objects are salient to the animals. Without isolating individual features, whole-object preference testing cannot diagnose the source of any observed bias, guide the design of improved alternatives, or determine whether validation transfers across species.

If species ecology predicts which features are salient, the consequences follow directly. Validation cannot be species-neutral; features that align with a species’ natural behavioral repertoire should be salient, while perceptual differences less related to that repertoire should be neutral. We tested this prediction directly using a paired-object design in which objects within each pair differed along exactly one researcher-defined feature, allowing assessment of whether that feature is salient to the animal. We applied the protocol to two wild-caught mouse species with contrasting ecologies. Wood mice (*Sylvaemus uralensis*, Pallas, 1811) inhabit mixed and deciduous forests in Central Eurasia (Gromov & Erbaeva, 1995)[9]; comparative studies describe *S. uralensis* as arboreal, a good climber and jumper that forages in trees and may nest above ground (Wilson *et al.*, 2017)[10], with postcranial skeletal proportions consistent with climbing activity in forest-dwelling *Sylvaemus* species (Kuncová & Frynta, 2009)[11]. In contrast, striped field mice (*Apodemus agrarius*, Pallas, 1771) are described as terrestrial, with climbing activity moderate compared with that of congeners (Wilson *et al.*, 2017)[10], occupying open habitats such as meadows and floodplains across Eurasia (Gromov & Erbaeva, 1995)[9]. Wild-caught rodent species differ in general exploratory tendency under controlled laboratory conditions (Galsworthy *et al.*, 2005)[12]; whether they also differ in which object features they treat as salient is the question we address here.

We predicted that if height differences are ecologically relevant, *S. uralensis* should respond differently to height-varying objects compared to *A. agrarius*, and that this difference should be consistent with their respective behavioral repertoires. Specifically, we expected *S. uralensis* to prefer taller objects or avoid shorter ones, reflecting their arboreal tendencies. The height prediction was comparative: it followed from the contrast between the two species’ ecologies and specified which species should respond and how. For two further features we held exploratory predictions – that eye-like motifs might be aversive, or that apertures might attract investigation – but these rested on general ecological reasoning applying to both species, and entailed no directional prediction about a species difference. Other pairs served primarily to assess the baseline level of feature salience.

## Materials and Methods

### Animals

The sample comprised 17 male mice of *Sylvaemus uralensis* (8 adults, 9 subadults) and 14 male mice of *Apodemus agrarius* (11 adults, 3 subadults). Animals were live-trapped in their natural habitats using Sherman-type live traps. *S. uralensis* were captured in a mixed forest and *A. agrarius* in a floodplain meadow on the bank of the Moscow River, both at the Zvenigorod Biological Station named after Skadovsky (Moscow Region, Russia). Age class (adult or subadult) was determined by anogenital distance and body size. After capture, animals underwent a minimum one-week habituation period. They were housed in groups of 2–5 animals in standard laboratory cages with food and water available ad libitum under a natural light cycle (approximately 04:30– 20:30 MSK). All procedures were approved by the Moscow State University Bioethics Commission (meeting No. 156-d, 16.11.2023, application No. 165-a).

### Experimental Arena

A rectangular arena with opaque white walls and rounded corners (80 × 40 cm floor; 59.5 cm wall height) was used. The floor was sheet vinyl divided into zones: two circular object sectors (diameter 20 cm), a central corridor, and a 10 cm wall-adjacent border (Fig. 1A). Video was recorded at 25 frames per second from a fixed camera (Sony Digital, Sony Corporation, Tokyo, Japan) positioned 160 cm above the arena. Illumination was provided by two 40 W incandescent diffused-light bulbs positioned 40 cm apart at camera height.

**Figure 1.**
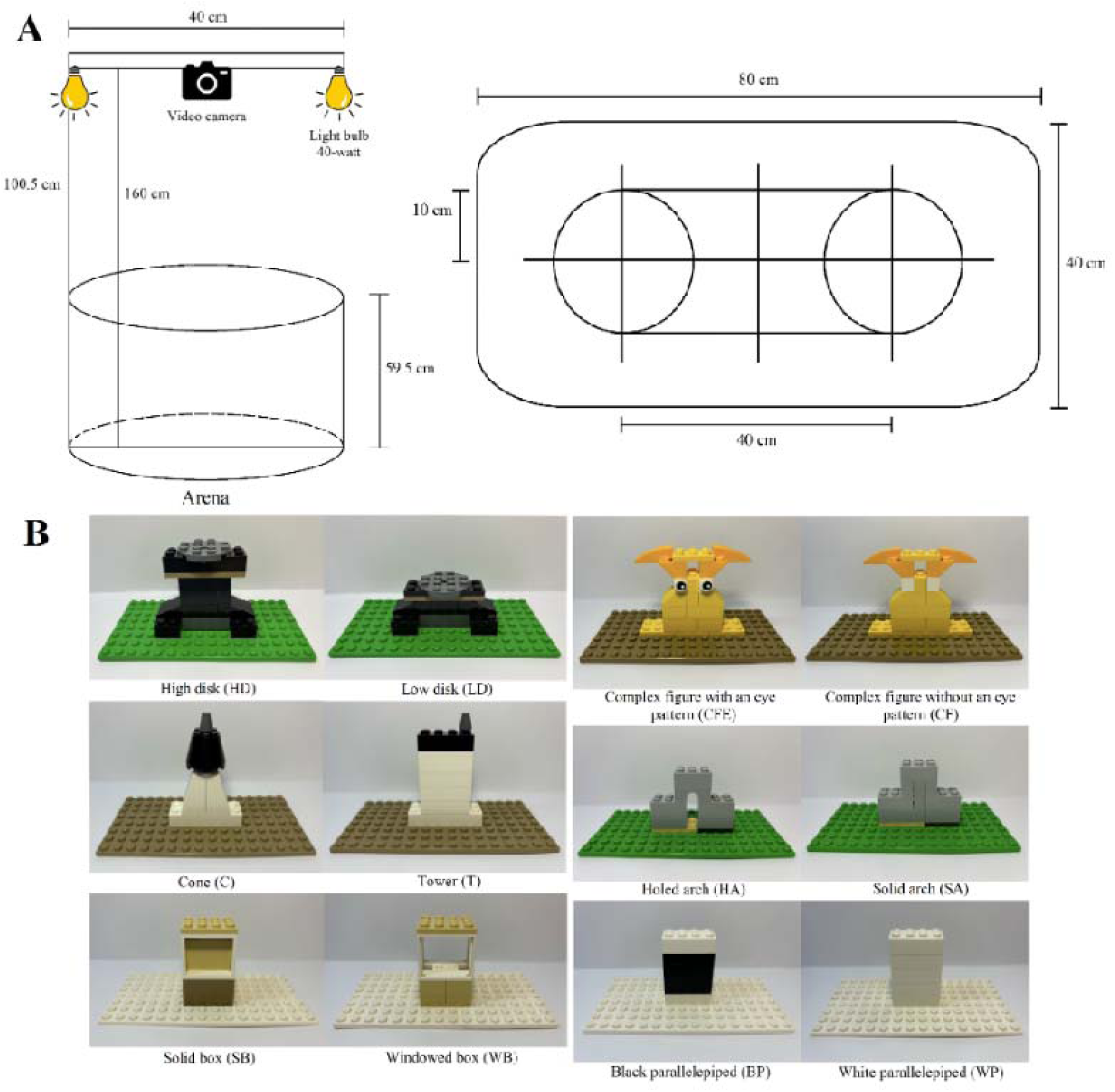
**A**. The experimental arena and setup. **B.** Pairs of novel standardized objects used in the experiment.

### Objects

Six pairs of objects were assembled from Lego components (LEGO Group, Billund, Denmark). Within each pair, objects differed along exactly one feature (Fig. 1B, Table 1): shape of the apex (pointed vs. flat top), stand height (high vs. low), presence of an aperture (holed vs. solid), color in the center (white vs. black), or presence of an eye-pattern motif. Each object was mounted on a Lego base plate; base plate color was identical within pairs. In pairs where the feature is aperture presence (pairs 3 and 5), weight differences are unavoidable as material removal reduces mass (2.0 g and 6.4 g differences, respectively). Pair 1 also has a 5.2 g weight difference and Pair 2 a 6.6 g difference. These differences may be perceptible through tactile manipulation and represent a potential secondary source of variation. All objects rested on wide base plates that provided stable footing but were not fixed to the arena floor. When an animal dislodged an object by climbing, far more frequently in *S. uralensis* than in *A. agrarius*, the experimenter returned it to the zone with the trial clock running.

**Figure 2.**
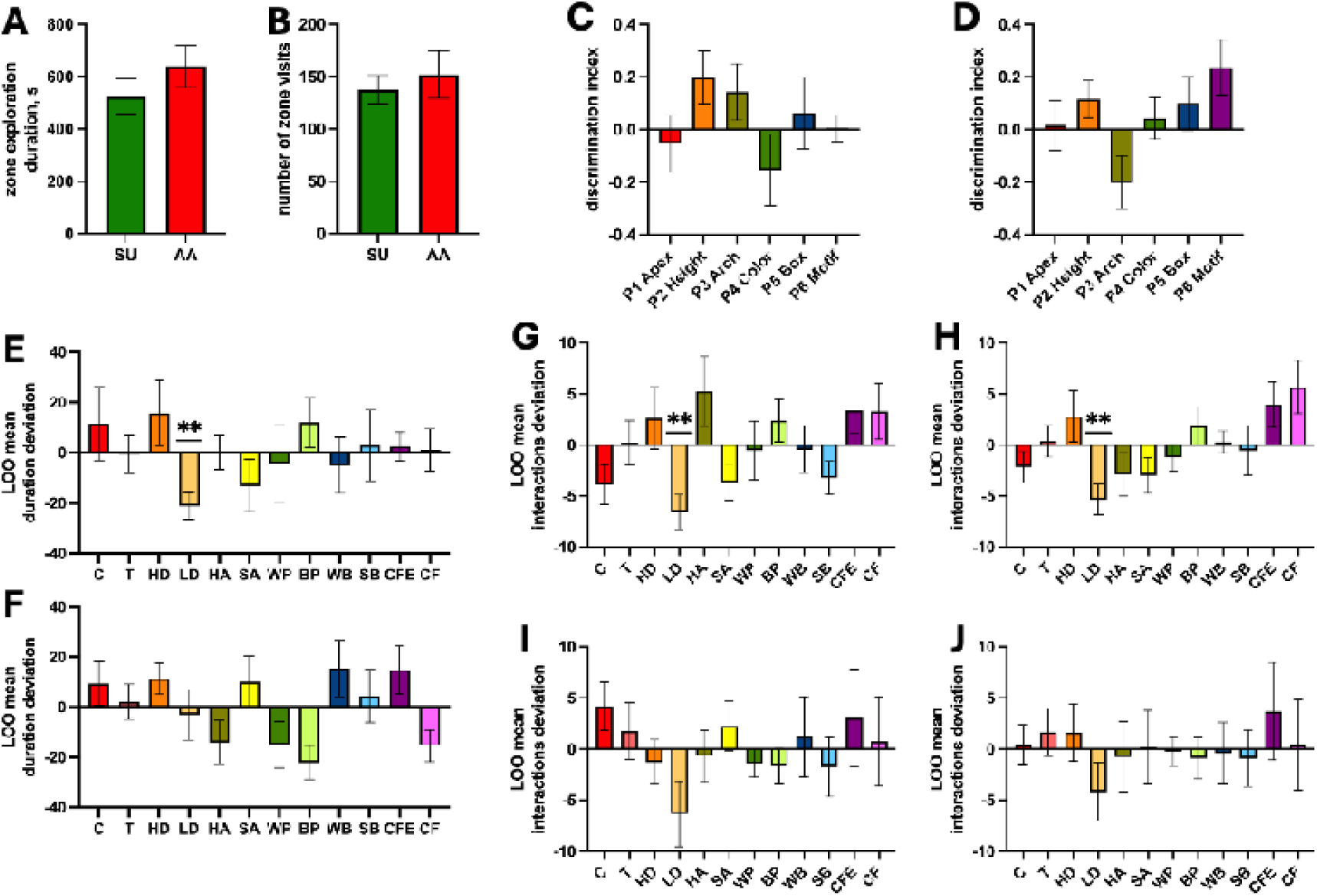
Species-wise behavioral comparisons. SU, *Sylvaemus uralensis*; AA, *Apodemus agrarius*. Values are mean ± s.e.m. **(A,B)** Interspecies comparison of aggregate object zone visit duration and number of visits (Welch’s t-test). **(C,D)** Within-pair discrimination indices for zone visit duration by species (planned comparisons against zero with Šídák correction for 6 comparisons) (C: SU; D: AA). **(E,F)** Leave-one-out deviation scores for object zone visit duration by species: each bar shows the mean deviation of that object’s duration from the animal’s mean across all other objects (one-sample t-test against zero, Šídák-corrected for 12 comparisons) (E: SU; F: AA). **(G,H)** Leave-one-out deviation scores for interactions in *S. uralensis* (G: touches; H: climbs). **(I,J)** Leave-one-out deviation scores for interactions in *A. agrarius* (I: touches; J: climbs). ** – p < 0.05 after Šídák correction.

**Table 1.**
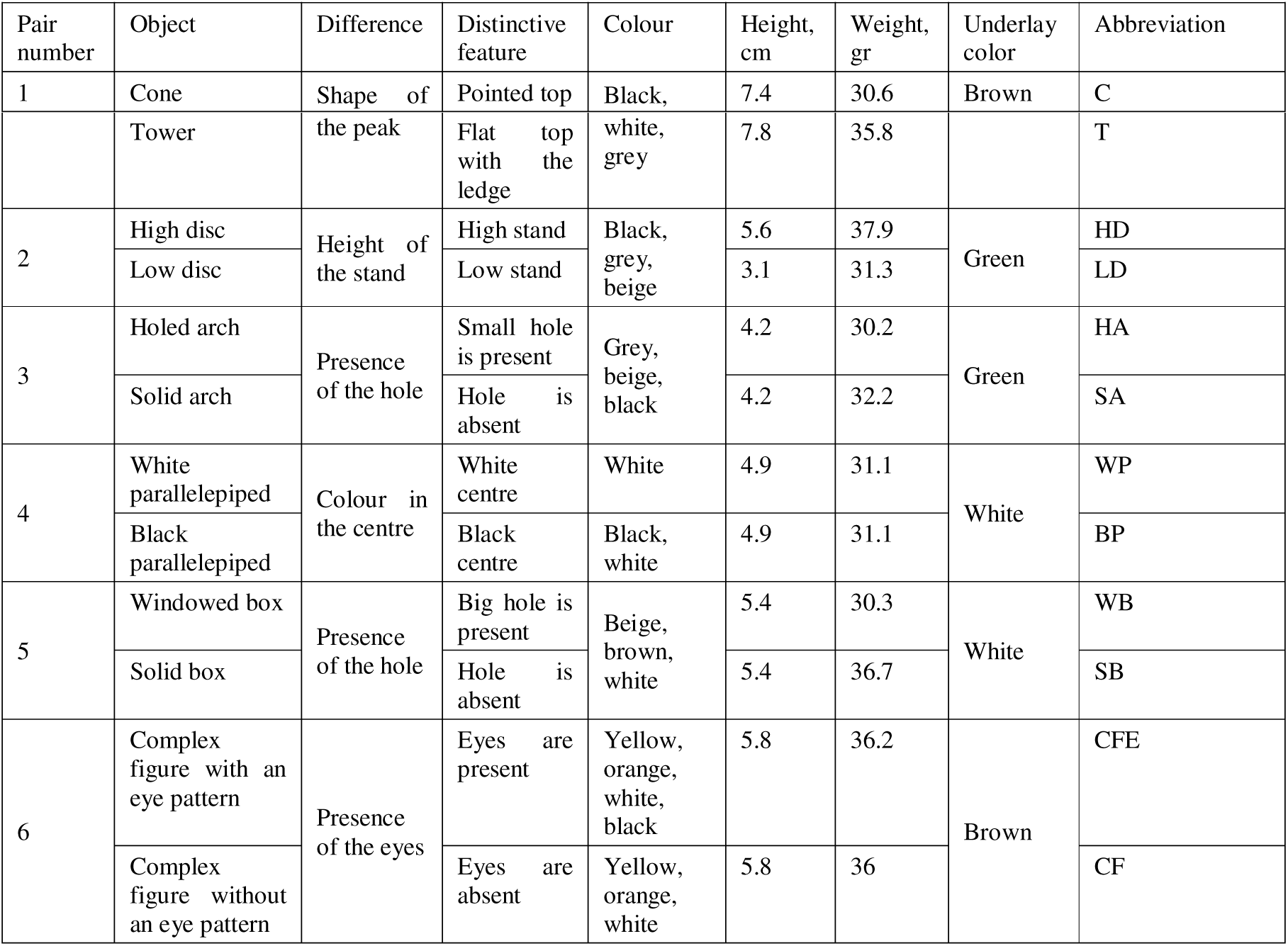
Pairs of experimental objects.

### Procedure

Day 1: 5-minute habituation to empty arena. Days 2–4: object exploration trials. Each pair was presented once to each animal (six trials total per animal). Trials lasted 8 minutes with 5-minute inter-trial intervals. Half the animals were tested in morning sessions (09:00–13:00), half in evening (18:00–22:00), assigned pseudo-randomly. Object pairs were assigned to trials in pseudo-random order. One object was placed in each object sector, 40 cm apart measured from object centers. Arena and objects were cleaned with water then 55% ethanol after each animal. Objects were taken off the base plate and both objects and plates were submerged in 55% ethanol solution for 5 minutes and then dried with paper towels. Between trials of the same animal, floor ethanol cleaning was omitted. Two *S. uralensis* individuals (SU-4, SU-12) received pair 1 (apex shape) twice due to a counterbalancing error and were not presented pair 4 (color); their pair 1 data were excluded from all analyses, but data from unaffected pairs were retained.

### Behavioural Coding and Analysis

All videos were scored frame-by-frame by two observers (S.E.A., E.N.A.) who coded jointly through consensus discussion, using VLC media player (VideoLAN, Paris, France). The primary measures were the total duration spent in each object zone and the number of separate visits to each zone. Additional measures included tactile interactions (touching, biting, handling) and on-object behaviors (climbing, jumping onto, rearing upon). We note that zone presence includes both exploration and passive behaviors such as grooming. Raw data were recorded in Microsoft Excel 2013 (Microsoft Corporation, Redmond, WA, USA). Trials in which total object zone duration was less than 20 s were excluded, as this threshold represents the minimum exploration criterion for reliable behavioral assessment in object-based tasks (Leger *et al.*, 2013; Lueptow, 2017). Because our primary measure is zone presence rather than active nose-directed exploration, this criterion is conservative relative to the standard NOR threshold. Twelve trials (of 186 total) met this exclusion criterion. Sample sizes per pair after all exclusions ranged from 12 to 16 animals for *S. uralensis* and from 10 to 14 animals for *A. agrarius* (see Supplementary Table 2 for details). Age class (adult vs. subadult) did not significantly affect discrimination index in *S. uralensis* (unpaired t-tests, all p > 0.19). The *A. agrarius* sample was too unbalanced (11 adults, 3 subadults) for meaningful age comparison. Age was not included as a covariate in subsequent analyses. Discrimination index (DI) was calculated as: DI = (Time_a − Time_b) / (Time_a + Time_b). DI ranges from +1 (exclusive preference for object A) through 0 (equal) to −1 (exclusive preference for object B). The primary analysis used a two-way mixed-effects model (REML estimation) on discrimination indices, with Species (*S. uralensis* vs. *A. agrarius*) as a between-subjects factor and Pair (six levels) as a within-subjects factor. Planned comparisons tested each pair’s DI against zero within each species using one-sample t-tests with Šídák correction for six comparisons. To assess individual object effects without the circularity inherent in comparing objects to a grand mean that includes them, we computed leave-one-out (LOO) deviation scores: for each animal and each object, the deviation was calculated as that object’s zone duration minus the mean duration across all other objects for that animal. LOO scores were tested against zero using one-sample t-tests with Šídák correction for 12 comparisons. The same LOO approach was applied to interaction counts (total interactions, touches, and climbing). Weight confound analyses tested whether the heavier object in weight-discrepant pairs (1, 2, 3, 5) was systematically preferred, using one-sample t-tests of weight-directed DI against zero with Šídák correction for four comparisons. Analyses were conducted in GraphPad Prism 9.5.1 (GraphPad Software, San Diego, CA, USA).

## Results

### Overall exploration

Total object zone visit duration and number of visits did not differ between species (Welch’s t-test; duration: p = 0.27; visits: p = 0.60). Both species showed comparable exploratory motivation, providing a baseline against which object-specific effects can be interpreted.

### Within-pair discrimination indices

The mixed-effects model on duration discrimination indices revealed no significant main effect of Pair (F(5, 122) = 1.532, p = 0.18), no main effect of Species (F(1, 29) = 0.091, p = 0.76), and a trending but non-significant Pair × Species interaction (F(5, 122) = 2.220, p = 0.056). The mixed-effects model on visit frequency DIs yielded similar results (Pair: F(5, 122) = 1.724, p = 0.13; Species: F(1, 29) = 0.014 p = 0.91; interaction: F(5, 122) = 1.717, p = 0.13). Planned comparisons of individual pair DIs against zero confirmed that no pair produced a statistically significant preference in either species after Šídák correction (all corrected p > 0.13). This pattern held for pairs differing in color (pair 4), aperture presence (pairs 3 and 5), apex shape (pair 1), and decorative motifs (pair 6).

### Leave-one-out analysis

Although within-pair DI for pair 2 (height: high disc vs. low disc) did not reach significance after correction in either species (*S. uralensis*: mean DI = +0.20, raw p = 0.066; *A. agrarius*: mean DI = +0.12, raw p = 0.13), the leave-one-out analysis revealed a clear species-specific pattern at the individual object level. *S. uralensis* showed significant avoidance of the low disc (LOO deviation = −21.10 s, t(15) = 4.032, raw p = 0.0011, Šídák-corrected p = 0.012). This was the only object to deviate significantly from baseline in either species after correction for 12 comparisons. The avoidance was corroborated by interaction data: wood mice touched the low disc less than their personal average (LOO touches: t(15) = 3.818, corrected p = 0.018) and climbed it less (LOO climbing: t(15) = 3.591, corrected p = 0.029). No other object showed significant deviation on any interaction measure in this species. *A. agrarius* showed no significant deviation for any object on any measure after correction (all corrected p > 0.06). In particular, the low disc deviation was near zero (−3.06 s, p = 0.99), indicating no comparable avoidance in this species.

### Visit frequency

Visit frequency DI did not differ significantly from zero for any pair in either species (all corrected p > 0.19), including pair 2 (height), where the duration DI showed a nonsignificant trend.

### Weight controls

For pairs with weight confounds (pair 1: Δ = 5.2 g; pair 2: Δ = 6.6 g; pair 3: Δ = 2.0 g; pair 5: Δ = 6.4 g), discrimination indices oriented toward the heavier object did not differ from zero in either species (all Šídák-corrected p > 0.13).

## Discussion

### Ecology and feature salience

The central finding of this study is that ecological background and behavioral repertoire predicted the response to one object feature, height, in a species-specific way. Height difference was salient to *S. uralensis*: they showed systematic avoidance of the shortest object, expressed consistently across zone duration, touching, and climbing. This pattern matches a species whose habitat contains prominent vertical structure and whose behavioral repertoire includes regular vertical locomotion (Kuncová & Frynta, 2009; Wilson *et al.*, 2017)[10,11]. *A. agrarius*, which occupy more open habitats where vertical structure is less common (Gromov & Erbaeva, 1995)[9], showed no equivalent avoidance. The convergence across three independent measures strengthens the inference that the effect reflects differential engagement with the object rather than a statistical artifact, and the species difference is in the direction predicted by the ecological backgrounds of the two species.

Specifically, *S. uralensis* in our data avoided the low disc rather than preferring the high one, a reading supported by the significant negative LOO deviation for the low disc in the absence of a corresponding positive deviation for the high disc. Functionally, this suggests that the low disc was treated as an inadequate structure, one that fails to afford the climbing or vertical locomotion opportunities that forest-dwelling *S. uralensis* routinely exploit (Wilson *et al.*, 2017)[10]. Under this interpretation, objects below a certain height may be functionally equivalent to flat substrate and therefore less ecologically relevant than taller alternatives. We cannot, however, from two species sampled from single populations, claim that ecology determines the salience of the object features we employed in this study as a generally applicable rule, and while captive rearing alone appears not to reshape wild behavioural repertoires (Künzl *et al.*, 2003)[13], domestication does, so these findings may not transfer to laboratory strains.

### The null pairs and the role of behavioral repertoire

The null results in the other five pairs are informative in their own right. Across both species and after correction, neither colour, apertures, apex shape, nor decorative motifs produced within-pair preference. One possibility is that wild-caught rodents discriminate between features that intersect with their species-typical behavioral repertoire without prior training, and respond less differentially to features that fall outside it. The broader principle that animals prioritize ecologically meaningful stimulus dimensions over arbitrary perceptual ones is well established in the comparative cognition literature (Shettleworth, 2010)[1]. Lego objects do not carry traces of conspecific or predator activity, and their colour and apex-shape variations have no evident connection to the behavioral repertoires of either species.

Height, by contrast, is a physical property directly relevant to species with the corresponding repertoire and less so to those without it. Two of the null features, however, were not arbitrary by our own account. We expected apertures to attract investigation as potential refugia, and eye-like motifs to be aversive as predator cues. Both predictions rested on general ecological reasoning, but not on the ecological contrast between the two species: refuge and predator cues are relevant to each, and neither prediction specified a species difference. Only the prediction that did follow from the species contrast, height, held. This bears on how features might be chosen for validation. Ecological reasoning about a species’ general biology did not identify which features would prove salient, and we could not have told in advance which of our three predictions would survive testing. What settles the question is systematic pairwise testing.

### Methodological observations

Several features of our analysis may affect how object-based preference studies might be designed and analyzed.

The height effect in *S. uralensis* was detectable through the leave-one-out analysis but not through within-pair discrimination indices, which were null for every pair in both species after correction. The two approaches ask different questions: a within-pair index compares an object only against its partner, whereas the leave-one-out deviation compares each object against the animal’s own exploration baseline across all other objects. An effect can therefore surface on one index and not the other. The height response appeared as a departure from baseline but not as a within-pair preference, which is why the leave-one-out analysis registered it and the pairwise index did not, a fact about the form the effect took, not about one test being the better instrument.

Across our data, zone occupancy duration was more informative than the number of zone visits: visit-frequency indices did not differ from zero for any pair in either species, including the height pair where duration carried the effect. Where feature-driven preferences exist, they appear to be expressed as differences in how long an animal stays rather than how often it approaches – a pattern worth minding when selecting dependent measures for object-based tasks.

A related point concerns the resolution of the behavioral readout. Our primary measure, zone occupancy, is coarse: time in an object’s zone includes investigation but also grooming, resting, and passage, and on its own it cannot distinguish engagement with the object from mere presence near it. The object-directed interaction measures, touching– and climbing-related behaviour, do not have this ambiguity. The low-disc effect appeared on all three – zone duration, touches, and climbing – so the avoidance reflects reduced engagement with the object, not just less time spent in its part of the arena.

The height pair was also weight-discrepant, the low disc being lighter as well as shorter. Because no other weight-discrepant pair showed a systematic preference for the heavier object (Results), we attribute the height effect to height rather than mass, while noting that touching and climbing cannot separate the two within that pair.

Our reliance on modular construction elements was itself a standardization choice. Assembling objects from a common component system let us hold all dimensions constant except the single manipulated feature, and makes the exact objects reproducible by any laboratory with the same components – the same reproducibility goal pursued through 3D-printed object libraries (Inayat *et al.*, 2021)[8], reached by different means. The two approaches are complementary: physical reproducibility fixes the standardization problem, while paired-feature testing establishes which of the objects’ features are salient to a given species.

### Limitations

Several features of the design constrain interpretation. First, the age structure of the sample: both species samples mixed adults and subadults. Age class did not affect the discrimination index in *S. uralensis* (all *p* > 0.19), but the *A. agrarius* sample was too unbalanced (11 adults, 3 subadults) to test the same question, so we cannot formally exclude an age contribution to the species difference, although the evidence available in *S. uralensis* argues against one.

Second, the coding procedure. Videos were scored by two observers working jointly through consensus rather than independently. Consensus scoring reduces the idiosyncratic error a single coder can introduce, but because the two observers did not score independently, we cannot provide a quantitative inter-observer reliability estimate.

Third, cumulative exposure. Each animal encountered all twelve objects across the six trials. The within-subjects design was deliberate: it draws the most information from a modest wild-caught sample and lets each animal serve as its own baseline, which the leave-one-out analysis requires, since a deviation score is defined only once an animal has met the full object set. The cost is that later trials build on accumulating experience with the objects as a whole; presenting pairs in pseudo-randomized order spreads any such effect across objects rather than loading it onto particular ones, but the design does not separate first-exposure from later responses.

Fourth, object displacement. Objects were occasionally dislodged and repositioned during trials; the low disc was displaced least frequently of all twelve objects, so this intervention works against the reported effect rather than producing it.

Fifth, our animals came from single populations of each species; behavioural profiles vary between populations (Dammhahn *et al.*, 2020)[14], so generalization even within species is uncertain, consistent with broader concerns about sample representativeness (Webster and Rutz, 2020)[15].

### Practical implications

Our observations point to a few concrete practices for studies that select or validate objects for recognition and exploration tasks. The most basic is to validate objects pairwise, varying one feature at a time: the informative outcome is not “equal preference” but the absence of a detectable within-pair preference, and when that coincides with no deviation from baseline on the leave-one-out metric, the two together set a ceiling that supports varying the feature freely thereafter. Which features warrant this scrutiny should be read from the study species’ ecology and behavioural repertoire, since features that intersect that repertoire, like height for an arboreal species or entry points for a fossorial one, are both the most likely to bias exploration and the most in need of validation. Crucially, that validation cannot be done by analogy: a feature neutral in one species, strain, or population cannot be assumed neutral in another, and should be checked in the animals actually to be used.

The underreporting documented by Koivisto *et al.* (2025)[5] is worth addressing directly, by reporting object dimensions, colour, material, and weight, and by stating which features were deliberately varied, whether they were tested for salience, and with what result. The pairing of analyses and of measures discussed above applies here too: both within-pair and leave-one-out comparisons, and both zone occupancy and object-directed interaction, are relevant in such a validation.

### Future directions

The account advanced here, that a species’ ecology and behavioural repertoire predict which object features are salient, generates predictions the same paired-object protocol can test. Texture is a tractable next dimension, since rodents discriminate it finely through the whisker system (Wu *et al.*, 2013)[16] and surface texture can shape preference and locomotion (Yoshida *et al.*, 2025)[17]. And comparing wild-caught animals with laboratory-reared conspecifics on identical pairs would test whether ecologically grounded salience survives the behavioural shifts of captive rearing – the question our wild-caught sample raises but cannot answer.

## Acknowledgements

We thank the staff of the Zvenigorod Biological Station for logistical support during animal capture.

## Declaration of Competing Interests

The authors declare no competing interests.

## Author Contributions

Conceptualization: M.G.P.; Methodology: M.G.P., A.M.Y.; Investigation: A.M.Y., E.A.S., N.A.E.; Formal analysis: A.M.Y., E.A.S., N.A.E.; Resources: V.M.M., V.Yu.O.; Writing – original draft: A.M.Y., E.A.S., N.A.E.; Writing – review and editing: M.G.P., V.Yu.O.; Supervision: M.G.P.; Funding acquisition: M.G.P. All living authors have reviewed and approved the final manuscript. M.G. Pleskacheva, who conceived and supervised the study, and V.M. Malygin, who provided zoological expertise and resources, are listed posthumously.

## Funding

This work was supported by the Moscow State University Development Program, project No. 23-Ш03-02.

## Ethics

All procedures were approved by the Moscow State University Bioethics Commission (meeting No. 156-d, 16.11.2023, application No. 165-a).

## Data Availability

All raw behavioural data are available in Supplementary Table 1. Additional data are available from the corresponding author upon request.

## Notes

### Competing Interest Statement

The authors have declared no competing interest.

### Summary of Updates

This version has been substantially revised from the originally posted preprint. The empirical data, experimental design, and core statistical results are unchanged; the revisions concern interpretation, framing, and scope. The most significant change is to the interpretation of the null results. The previous version explained the absence of preference in five of six object pairs through an "ecological valence" framework, proposing that wild rodents evaluate objects by biological relevance (food, shelter, conspecific or predator signals) rather than by discriminating physical features. This version removes that framework. In its place, we note that two of the null features were ones we had ourselves predicted to be salient on ecological grounds, and that the failure of those predictions indicates ecological reasoning alone does not identify which features matter and the question must be settled by testing. This shifts the paper's secondary claim from a hypothesis about how animals evaluate objects to a more defensible observation about the limits of a priori feature selection. The scope of the central claim has been narrowed accordingly. Where the earlier version stated that ecological background predicts which features are salient in general, this version distinguishes the comparative height prediction (which was confirmed) from general ecological predictions (which were not), and bounds the generalizability of the findings to wild-derived populations. Other changes: the Results and Discussion sections, previously combined, are now separated; the characterization of the leave-one-out and within-pair analyses as differing in sensitivity has been corrected to reflect that they test different questions; and the reference list has been updated to reflect the revised interpretation. The title has been changed from "A paired-object protocol for validating feature salience in rodent exploration: evidence that ecology predicts which features matter" to reflect this reframing: the earlier title led with the validation method and treated the ecological finding as secondary, whereas the present version foregrounds the finding itself.

